# Topical Delivery of Elastic Liposomal Vesicles for Treatment of Middle and Inner Ear Disease

**DOI:** 10.1101/2022.06.02.494551

**Authors:** Raana Kashfi Sadabad, Anping Xia, Nesrine Benkafadar, Chrysovalantou Faniku, Diego Preciado, Stella Yang, Tulio A. Valdez

## Abstract

We present a topical drug delivery mechanism through the ear canal to the middle and inner ear using liposomal nanoparticles without disrupting the integrity of the tympanic membrane. The current delivery method provides a non-invasive,and safer alternative to trans-tympanic membrane injections, ear tubes followed by ear drops administration and systemic drug formulations. We investigate the capability of liposomal NPs, particularly transfersomes (TLipo), used as drug delivery vesicles to penetrate the tympanic membrane (TM) and round window membrane (RWM) with high affinity, specificity, and retention time. The TLipo is applied to the ear canal and found to pass through tympanic membrane quickly in 3 hours post drug administration. They are identified in the middle ear cavity 6 hours and in the inner ear 24 hours after drug administration. We performed cytotoxicity in vitro and ototoxicity in vivo studies. Cell viability shows no significant difference between the applied TLipo concentration and control. Furthermore, auditory brainstem response (ABR) reveals no hearing loss in 1 week and 1 month post administration. Immunohistochemistry results demonstrates no evidence of hair cell loss in the cochlea at 1 month following TLipo administration. Together, the data suggested that TLipo can be used as a vehicle for topical drug delivery to the middle ear and inner ear.

## Introduction

Middle ear diseases such as otitis media affect millions of people worldwide. Otitis media (OM), refers to a continuum of inflammatory conditions in the middle ear, including acute otitis media (AOM) and chronic middle ear effusion (COM) [1, 2]. OM is the second most common illness diagnosed in children in the US, with over eight million cases each year [3]. OM is also the most common cause for hearing impairment, surgery, and antibiotic prescription in pediatric patients [3, 4]. Over 95% of US children experience at least one episode of OM at some point in their childhood, 40% experience four or more episodes by the age of seven, and 26% require insertion of ventilation tubes [5]. Untreated OM can lead to complications such as cholesteatoma (a destructive middle ear lesion), speech and language development delays, and hearing loss [6, 7].

Topical drug delivery to the middle ear across the TM (i.e., eardrum) has gained increasing attention recently [8, 9], due to its advantages over the standard *systemic* drug administration methods such as oral intake or intravenous (IV) injection. While systemic methods can cause non-specific biodistribution which requires high-dosage administration, local delivery leads to a higher drug concentration at the site of infection without administering high quantities of medication while also reducing side effects and providing more overall convenience when it comes to treating children [9, 10].

The TM is a thin semi-translucent membrane comprised of three layers with an average thickness of 0.1 mm. The outer layer consists of stratified squamous keratinized epithelium and lipid-rich stratum corneum, with a similar structure to skin [11, 12], protecting the middle ear from objects entering the external auditory canal. Current methods of bypassing this barrier are to surgically insert a tympanostomy tube, intratympanic injection, or transtympanic diffusion, all of which have their limitations [9]. Surgical procedures such as the tympanostomy tube placement often requires general anesthesia and multiple post-operative check-ups until the tube extrudes and the tympanic membrane heals. Although in adults performing an intratympanic injection is possible, this is a painful procedure and has the risk of TM perforation. Polymeric hydrogels have been recently developed to reversibly enhance the influx of medicine through the TM using chemical permeation enhancers (CPEs) [11, 13].The latter approach is non-invasive, pain-free, and easier to apply to pediatric patients, and can offer targeted delivery to the middle ear infection site. However, combining multiple medications in a hydrogel solution is not easy when compared to nanoparticle formulations where each compound can be encapsulated separately.

More recently, nanoparticles (NPs) with carefully engineered structures and properties have proven effective for transdermal delivery (i.e., delivery through skin) for a range of cosmetic or therapeutic applications such as atopic dermatitis [14-16] and skin cancer treatment [17]. These carriers can penetrate into the skin without damaging it and deliver therapeutic agents loaded on or encapsulated within the NPs [16]. For ear applications, on the other hand, the effectiveness of NP-based delivery such as poly-lactic/glycolic acid (PLGA) has been demonstrated for delivery across the round window into the inner ear after intratympanic injections into the middle ear cavity. However, the effectiveness of NPs for transtympanic crossing – avoiding any incision of the TM whatsoever – and delivering the medicine to the middle and inner ear with high affinity, specificity, and retention time has not been demonstrated [18].

In this study, we use liposomal NPs [19], which are vesicular structures composed of phospholipids arranged in a shell-like bilayer with hydrophilic surface (facing aqueous solution) and hydrophobic interior. This shell-like structure makes liposomes ideal carriers, enabling loading or encapsulation of drug molecules. Their ability to fuse (at nanoscale) with lipid-rich barriers in skin or tympanic membrane enable liposomes to push the cargo (e.g., medicine) through without damaging those barriers. Moreover, liposomes are bio-compatible (i.e., non-toxic) and biodegradable, making them safe for administration to pediatric patients. We use a subclass of liposomes called transfersomes [20, 21], which are additionally equipped with surfactants such as sodium cholate, deoxycholate, span, tween, and dipotassium glycyrrhizinate. These surfactants make the lipid bilayers destabilized and increase the deformability of the NPs, thereby enabling them to squeeze through the channels that can be one-tenth smaller in diameter, in lipid-rich biological barriers such as the skin or TM. For example, Aly et al [22] have shown the importance of using liposomal vesicles to improve penetration of levofloxacin, a hydrophilic antibiotic, which is loaded into polyethylene glycol 400 (PEG 400), through the TM. Furthermore, transfersomes have shown promising results to carry ciprofloxacin, a common AOM antibiotic, through the TM with 26-76% drug loading capacity [23]. However, their penetration and affect beyond the middle ear and into the inner ear have not been reported so far.

We synthesize transfersomes by the lipid film hydration method [21] and labeled them with a Rhodamine-B (RhB) based fluorescent dye to verify their accumulation in the middle and inner ear by confocal imaging. We verified the vesicles’ size, shape, and stability using a particle size analyzer and cryo-electron microscopy, following freeze-drying of the vesicles and loading them with the fluorescence dye. We first verify the non-toxicity of the vesicles via in vitro experiments. We then study the penetration rate and distribution of the labeled vesicles into the middle and inner ear at different time points using confocal microscopy imaging. We show that in a short period of time (3 hours) the vesicles reach the middle ear in high concentrations and verify their presence in smaller amounts in the inner ear as well. Finally, we assess the safety of the vesicles for inner ear drug delivery, by recording the auditory function, visualizing the hair cells, and performing histological analysis.

## Experiments Materials

α-phosphatidylcholine from soybean (Avanti Polar Lipids, 99%), Lissamine™ Rhodamine B-1,2-Dihexadecanoyl-sn-Glycero-3-Phosphoethanolamine (Rhodamine DHPE,Thermo Fisher Scientific), Triethylammonium Salt, Sodium cholate hydrate (Sigma-Aldrich, ≥99%), Sucrose (Sigma-Aldrich), Phosphate buffer saline, ethanol, Chloroform and other chemicals and solvents were analytical grade and used as supplied.

### Characterization

Cryo-transmission electron microscopy (cryo-TEM) analysis was conducted using a TFS Glacios 200 kV with a Gatan K2 detector. The samples for Cryo-TEM observation were dispersed in ethanol and dropped on copper Quantifoil holey grids cupper grid and were plunge frozen on a Leica GP at 95% humidity with 2 μl sample at 20 degrees. Size and zeta potential were determined via dynamic light scattering (DLS) with a Malvern Zeta Sizer Nano ZS instrument (model ZEN 3600). All DLS measurements were done at 25°C, were repeated for 3 times for each sample, and finally the mean values were reported. An infinite microplate reader (Tecan, Durham, USA) was used to calculate cell viability at 450 nm (reference wavelength 650nm). A confocal laser scanning microscope (CLSM700, Zeiss) was used to image tissues using a 10X, 40X and 63X objectives. The Images were processed with ImageJ software.

### Synthesis of fluorescently labeled TLipo formulation

We prepared TLipo vesicles by rotary evaporation followed by an extrusion method. Soybean and sodium cholate at a ratio of (87:13 m/m) were dissolved in a mixture of chloroform/methanol (50/50 v/v) in a 50 mL flask. The mixture was rotated under vacuum by a rotary evaporation system at room temperature. During this process, the organic solvents were evaporated, leaving a thin film layer of lipid on the wall of the flask. The deposited film was hydrated by adding PBS buffer and stirred slowly at room temperature for 2h. We allowed the resulting suspension to swell overnight at 4°C, forming TLipo vesicles. We used an extruder system (Avanti Polar Lipids) to downsize the obtained vesicles through a sandwich of 100 nm polycarbonate membranes up to ten times. Lastly, we lyophilized the frozen TLipo by removing PBS/water under vacuum. We added sucrose during the freeze-drying process to protect the TLipo from undesirable aggregation/fusion of vesicles and to reduce the freezing stress.

For labeling the TLipo with the fluorescent dye, we used the same technique and added the fluorescent Lisaamine Rhodamine-B dye (to the mixture of the soybean and sodium cholate with the organic solvents. Then, the formed pink color thin film was hydrated with PBS and processed for the freeze-drying. We named the final pink powder product as TLipo-RhB and stored it at 4°C.

### In vitro cytotoxicity study

We performed a methyl thiazolyl tetrazolium (MTT) cell metabolic activity assay to evaluate the safety of the designed formulations for subsequent clinical trials. A number of 1 × 10^4^ Vocal fold fibroblasts (VFF) cells per well were seeded in RPMI media supplemented in a 96-well plate. The cells were incubated at 37°C with 5% CO_2_ for 24 h. The medium was removed and replaced with the fresh medium containing TLipo vesicles at different concentrations from 1.95 to1000 μg mL^−1^. The cells were incubated at 37°C for a further 24 h. The medium was removed and 100 μL per well of the MTT reagent dissolved in PBS incubated with the cells for 4 h. The MTT reagent was extracted carefully and replaced with dimethyl sulfoxide (DMSO) (100 μL per well). The absorption values of the wells were recorded at 540 nm using a microplate reader.

### Nanoparticle cell uptake study

We measured the cellular uptake of the fluorescently labeled vesicles (TLipo-RhB) in a macrophage Raw cell line (RAW264.7). The cells were seeded into an 8-well chamber slides at the density of 50000 cells/well with DMEM supplemented with 10% FBS. The cultures were maintained at 37°C and 5% CO2. After 24h, the cells were washed once with PBS and exposed to the DMEM solution containing the vesicles with concentration of 50 μg/mL for 4h. Replicated wells containing cells and cell culture medium with no particles were prepared as a negative control. After 4h, cells were washed three times with PBS to remove the free nanoparticles from the cell medium before imaging and fixed with 200 μL freshly made 4% formaldehyde in PBS for 15min. Following fixation, the cells were stained with DAPI and kept in the dark for 24h. The Images were captured using a LSM700 confocal microscope (Zeiss) at 10X, 40X and 63X magnification.

### In vivo animal studies

CBA/CAJ mice (4-6 weeks age) were used for tympanic membrane penetration studies. All animal studies were performed in full compliance with approved protocol by Institutional Animal Care and Use Committee (IACUC) at Stanford University (APLAC #33028). All procedures were performed under anesthesia using a mixture of anesthetic Ketamine/Xylazine agents. 1 month old CBA/CAJ mice were divided into five groups (n=5). These groups received treatment by depositing the formulation directly onto the ear canal (20mg/mL, 20 μL). We used each mouse’s left ear for the treatment and their right ear as control and received no treatment. For the middle ear delivery, the entire hearing apparatus including the eardrum, middle ear, and inner ear was harvested at 3h, 6h, 24h, 1-week, and 1-month post-treatment.

### Ex vivo evaluation of ear penetration and ear tissues immunostaining

To evaluate middle ear delivery, the middle and cochlea were harvested and fixed in 4% paraformaldehyde (Electron Microscopy Sciences, Hatfield, PA, USA) at 4°C overnight. Samples were then decalcified in 0.5 M EDTA (VWR, Radnor, PA) for 48 hours at 40°C, and washed three times in PBS. For cryosection preparation, middle ears and cochleae were immersed in a sucrose gradient (10%–30%) and embedded in OCT. Samples were collected in 10 μm sections. The sections were stained with DAPI (Abcam, #ab104139) for nucleus visualization in mounting medium and covered with a cover slip to visualize the living cell layers of the TM, middle ear and inner ear sections. Finally, we visualized the vesicle biodistribution throughout the ear using confocal microscopy (using fluorophore dyes to label the nanoparticles). For whole-mount preparations, the organ of Corti was dissected free of each cochlea under a stereo microscope.

### Auditory brainstem response (ABR)

The auditory function was assessed by recording auditory brainstem responses (ABR) to evaluate the long-term effects of our formulation. 1 month old CBA/CAJ mice were randomly divided into two groups and ABR responses were measured prior at 1 week and 1 month post treatment with TLipo vesicles. The ABR measurement were obtained from needle electrodes positioned at the bottom of the tympanic bulla and at the vertex of the head, with a ground electrode placed in the rear leg. A bioamplifier (DP-311, Warner Instruments, Hamden, CT, USA) was used to amplify the signal 10,000 times. Amplitude-intensity functions of the ABRs were obtained at each frequency tested (4, 6.3, 8, 10, 12.5, 16, 20, 25, 32 and 46 kHz) by varying the intensity of the tone bursts from 10 to 80 dB SPL, in 10 dB incremental steps. 260 responses at each sound level were sampled and averaged. The ABR threshold at each frequency was calculated with respect to three standard deviations above the background noise. After obtaining baseline measurement, 20 μl of TLipo was applied and the same measurement was repeated for all treated mice. The mice were sacrificed at each time point and the samples were collected for the toxicity evaluation. Recordings and analysis were performed blindly.

### Immunohistochemistry

For visualization organ of corti and the hair cells, we followed a procedure consisting of vibratome sectioning of the cochlea, immunohistochemical staining of the sections, followed by confocal imaging. The whole mount tissue was immersed in a blocking solution of 4% donkey serum (Millipore Sigma, Massachusetts, USA) and 0.1% Triton-X 100 in PBS (PBST) for 1 hour at room temperature and then incubated with the primary antibody (rabbit anti-myosin VIIa, 1:1000; Proteus Biosciences Inc., Ramona, CA) at 4°C overnight. Specimens were washed three times with PBST and then incubated with the secondary antibody (Alexa Fluor 546 donkey antirabbit, 1:500; Invitrogen, Grand Island, NY) at room temperature for 1 hour. After washing with PBS, specimens were mounted in anti-fade Fluorescence Mounting Medium (DAKO, Carpinteria, CA) and cover slipped. Images were captured using a LSM700 confocal microscope (Zeiss) at 10X magnification. The cryosections of middle ear and inner ear were washed with PBS 3 times and cover slipped. The images were also captured under LSM700 confocal microscope.

For analysis of hair cell patterns in the organ of corti, the otic capsule containing the membranous inner ear was dissected out by decapitation and opening of the skull at the midline. The whole inner ear was fixed in 4% PFA in PBS overnight and decalcified with EDTA solution (100mM) for three days. The samples were washed in PBS and embedded in 4% low-melting point agarose (BioRad# 1613101) and positioned the inner ear samples for vibratome sectioning in the appropriate plane under a microscope. The sections (80 μm) were washed with 1x PBS and Permeabilized in 0.5% Triton X in 1x PBS for 30 minutes at room temperature. Then the samples were incubated in a blocking buffer (1% BSA, 0.2% Triton X in 1x PBS) for 30 min at room temperature. Myosin 7A (a polyclonal rabbit antibody 1/1000, Proteus Biosciences Inc, #25-6790) and neuron-specific beta-III antibody (TuJ-1) were used to label hair cells and neurons. Chromatin was stained with DAPI (1/2000). The secondary antibody used was the anti-rabbit IgG conjugated to Alexa 594 (1/500, Thermo Fisher Scientific, #A-2120). The samples were washed with PBS and mounted on slides for confocal imaging using a LSM 700; Zeiss system. No fluorescent signal was detected in control specimens without primary antibodies.

### Histological assessments

For hematoxylin and eosin (H&E) staining, we randomly divided mice in two groups and treated them with the vesicles without any fluorophore dye on the left ear and no treatment on the right ear. We used one group to dissect out the middle ear and another group for the inner ear after 1 month of the treatment. The ear specimens were fixed, decalcified, and immediately dehydrated in the 70% ethanol at 4°C. We followed the protocol for H&E staining as described elsewhere. Briefly, on a tissue processor, we dehydrated the specimens in a series of ethanol solutions (70%, 95%, 100%, each 1 h), cleared the in xylene at 40°C, and then embedded with molten paraffin wax. We adjusted the middle ear and inner ear in a particular orientation during embedding to cut the desired sections and collected the horizontal sections. Finally, we placed the sections on the microscope slides and stained the paraffin sections with hematoxylin and eosin (H&E).

## Results and Discussion

### TLipo characterization

We used a Rhodamine-labeled glycerophosphoethanolamine lipid (Rhodamine-DHPE) with excitation/emission maxima of ∼560/580 nm to label the TLipo vesicles for in vivo study of tympanic membrane delivery. We measured and compared vesicle sizes and shapes, with and without the fluorescent labeling, and after freeze-drying. We visualized the samples under Cryo-TEM, verifying uniform and unilamellar formation of vesicles with apparent size of ∼120 nm as shown in Figure 1a. The Cryo-TEM images confirmed the vesicles stayed intact without aggregation after freeze-drying and labeling with the fluorescent RhB-dye. To investigate the size distribution, we used a nano sizer system and verified the uniform vesicles size distribution at ∼159 nm for unlabeled and ∼177 nm for labelled vesicles, as shown in Figure 1b. We visualized the incorporation of Rhodamine-DHPE dye into TLipo using UV-Vis spectrometry with RhB absorption peak at ∼560 nm. We obtained a loading capacity (LC%) of 15% for the RhB dye in the TLipo vesicles by plotting the intensity of RhB-TLipo absorbance against the concentration of free Rhodamine-DHPE dye at six different concentrations.

**Figure 1.**
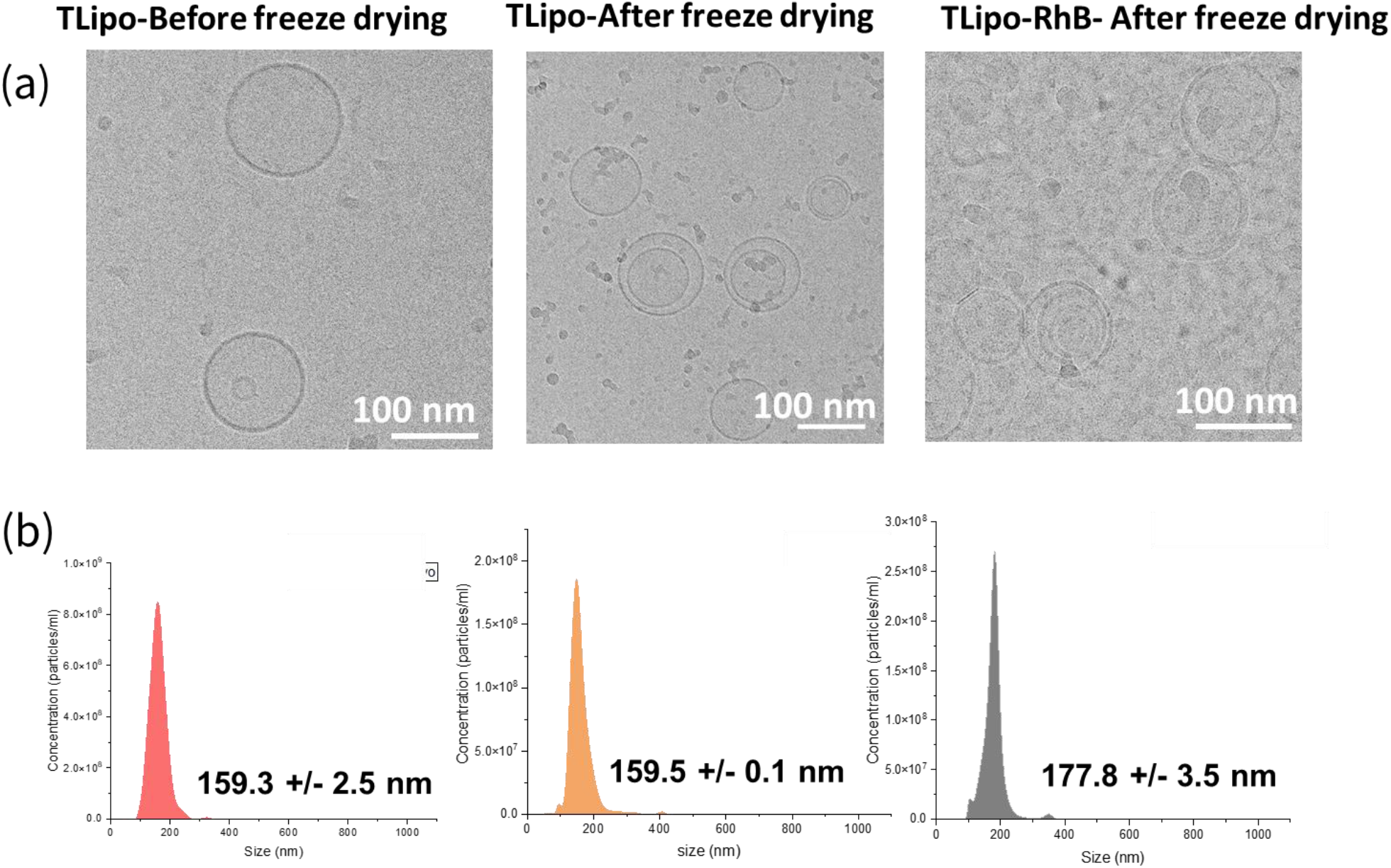
(a) Cryo-TEM images; (b) vesicles size distribution of TLipo vesicles before and after freeze-drying and encapsulating with RhB fluorescent dye.

### TLipo shows no evidence of cell toxicity

We studied the safety of TLipo vesicles by measuring the cell viability via a (3-(4,5-dimethylthiazol-2-yl)-2,5-diphenyl tetrazolium bromide (MTT) assay and measured the conversion of MTT in insoluble formazan crystals to assess mitochondrial metabolism. We treated vocal fold fibroblast (VFF) cells with TLipo vesicles at different concentrations for 24h at 37°C and the viability of cells were quantified by measuring their absorbance at 540 nm using a microplate reader. As shown in figure 2(a), the TLipo vesicles did not noticeably affect the proliferation of the cells, hence are not toxic.

**Figure 2.**
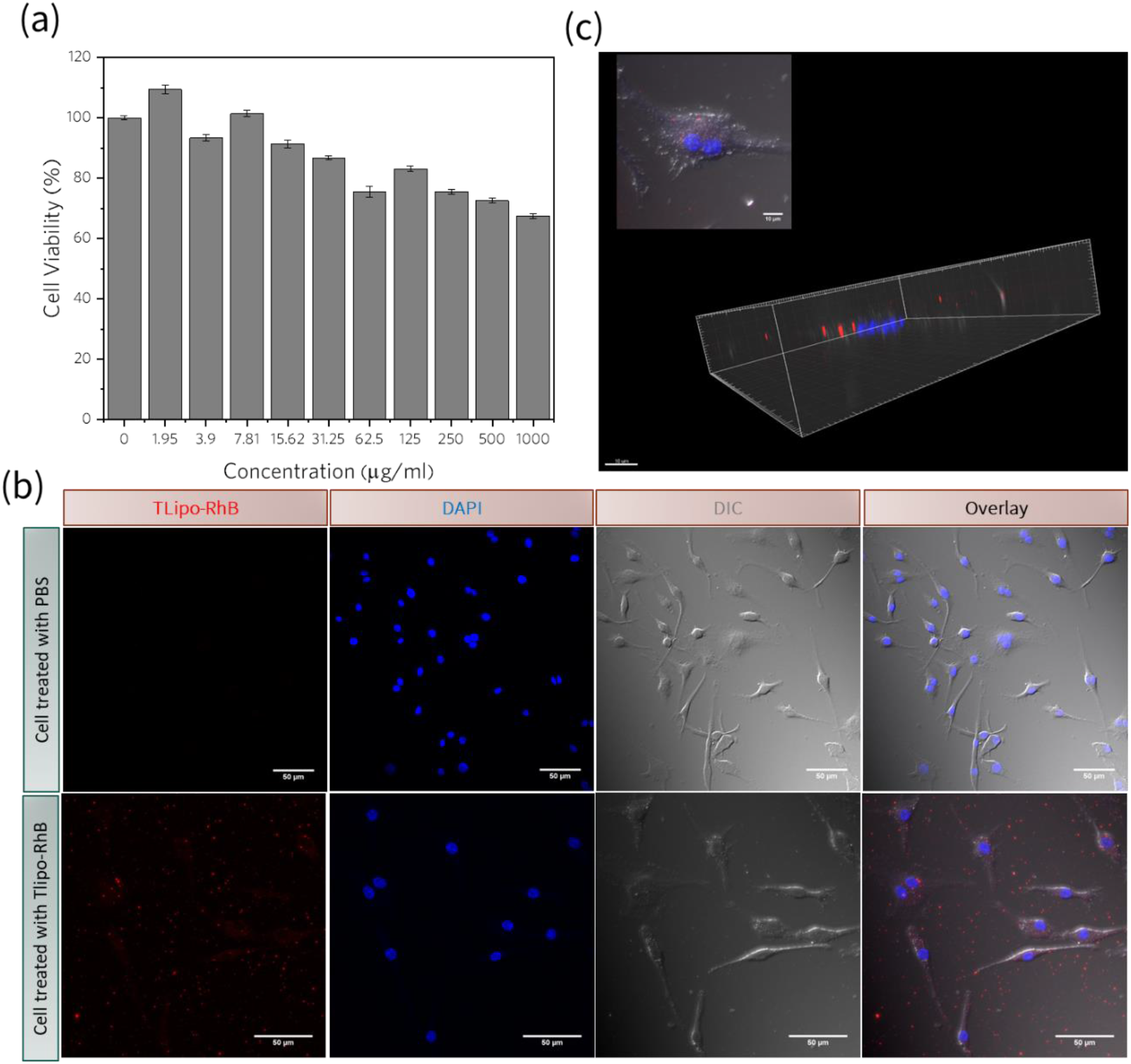
(a) MTT cytotoxicity studies on vocal cells after 24 h of incubation with TLipo vesicles (b) in vitro cellular uptake of Tlipo-RhB vesicles in macrophages at 4h post-incubation; (c) representative confocal images at single cell resolution analyzed by Imaris. The cell nuclei are stained in blue using the DNA-binding dye DAPI while red spots represent TLipo-RhB vesicles. Confocal microscopy images confirm the uptake of Tlipo-RhB by macrophages and MTT shows TLipo are not toxic.

We evaluated cellular uptake of the RhB-TLipo vesicles by macrophages. Macrophages infiltrate the middle ear during OM and are the first cells to invade the tissue after OM induction [24]. Macrophages resist penetration by many drugs and antibiotics [25]. We treated the cells with 50 ug/mL of the RhB-TLipo and subjected them to confocal microscopy (Figure 2b). We verified the penetration of the vesicles into the macrophages, to use them as potential carriers of the antimicrobial drugs in future studies, especially for acute OM cases with bacterial infections. We looked at the macrophages under confocal scanning microscopy after 4h incubation and localized their internalization by analyzing confocal images. Figure 2c shows the 3D cell visualization using the Imaris software (http://bitplane.com).

### TLipo crosses the TM into middle ear

After verifying the safety of the TLipo-RhB vesicles, we administered them as ear drops into the left ear of healthy mice without any sign of infection in their middle ear or TM perforation. We used the untreated right ear as the control case for all studies. We collected the ear samples at specific time points and processed them for the confocal microscopy imaging. We orientated and cut the ear samples under the microscope in a direction that gave us a better view of the TMs and middle ear areas. As shown in Figure 3, the ear sections consist of three major parts: the external ear canal, the TM, and the middle ear. The TM separates the middle ear from the external ear canal and is shown as a thin line in all images. The red staining in the images indicates TLipo-RhB distribution and location in the mice ear after 3h, 6h, 1 week and 1 month post treatment. For comparison, we tested the RhB dye without vesicles after 6h post-treatment. The mice without any treatment have been shown as control in Figure 3. We observed rapid distribution of TLipo-RhB after 3h through and beyond the TM in the middle ear cavity while no fluorescence was observed for the RhB dye after 6h post-treatment (n=5). These observations show the significant role of the TLipo vesicles in carrying the dye through the TM layers into the middle ear. Higher amounts of the fluorescent TLipo-RhB were observed in the middle ear 6h post-treatment, showing higher accumulation of the vesicles in the middle ear over time. After 1 week, the fluorescent TLipo-RhB in the middle ear start decaying and completely disappear after 1 month. This could be explained by vesicles degradation and clearance from the ear environment.

**Figure 3.**
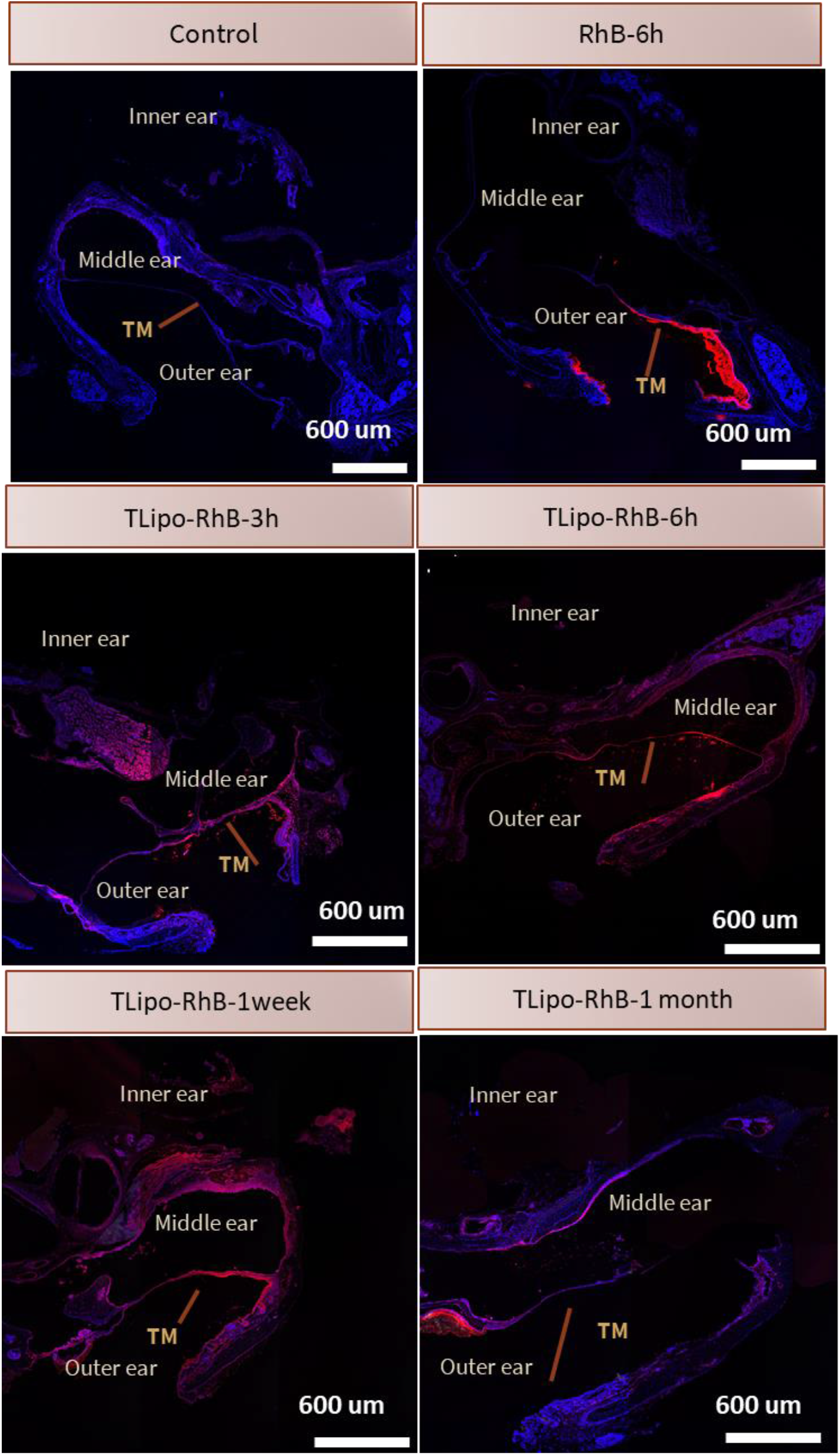
Confocal images of eardrum and middle ear for control (only PBS buffer), RhB, and TLipo-RhB penetration at different time points (n=5).

We looked at the TM at a higher magnification under confocal microscopy for the same ear samples treated with TLipo-RhB, RhB, and control after 6h post-treatment. We observed the structural layers of the TM and vesicles as shown in Figure 4 and verified the presence of TLipo-RhB at different layers of the TM with the ability to penetrate into the middle ear cavity. Whereas RhB solution without vesicles remained on the outer part of the TM, without penetration to the middle ear. No fluorescent signal was observed from the middle ear cavity for the free RhB.

**Figure 4.**
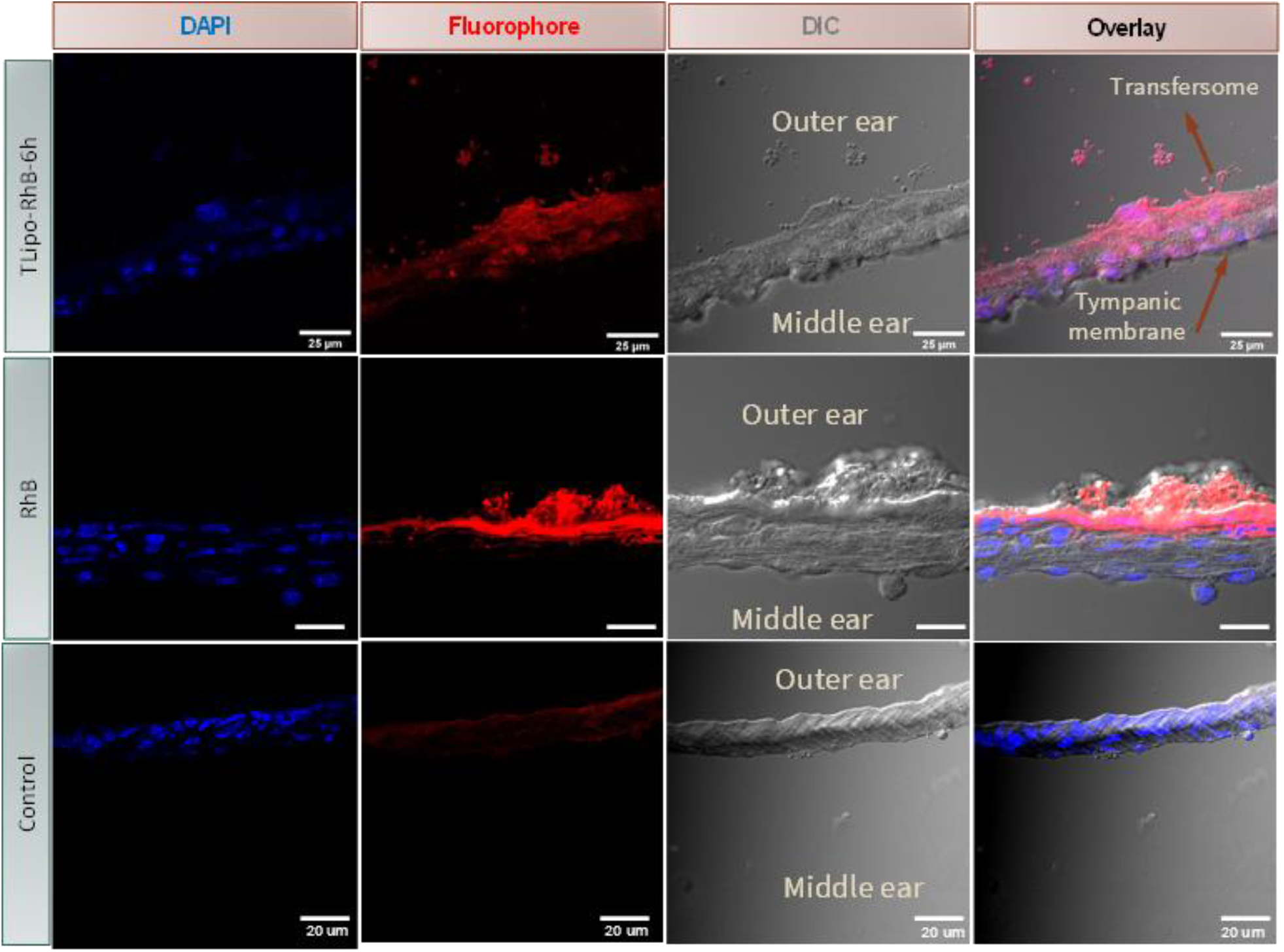
High magnification confocal microscopy of eardrum for control (only PBS buffer), RhB, and TLipo-RhB penetration after 6h of drop administration. The red color represents free RhB fluorescent dye and RhB encapsulated TLipo vesicles (n=5).

### TLipo enters the cochlea

After observing rapid diffusion of the vesicles through the TM into the middle ear, we next investigated penetration of the vesicles into the cochlea through the TM, using ex vivo imaging via confocal microscopy (Figure 5). The inner ear is more challenging to access by drugs and is located in one of the densest bones in the body called the otic capsule. There are several other barriers such as the round window and the oval window which limit local drug delivery to the inner ear through the middle ear route. To examine penetration into the inner ear, we deposited the TLipo-RhB vesicles (20mg/ml, 20 μl) on the eardrum and collected the mice cochlea 24h post-treatment. We were able to spot the vesicles at several locations in the treated inner ear, but none in the control ear as shown in Figure 6 (enlarged images).

**Figure 5.**
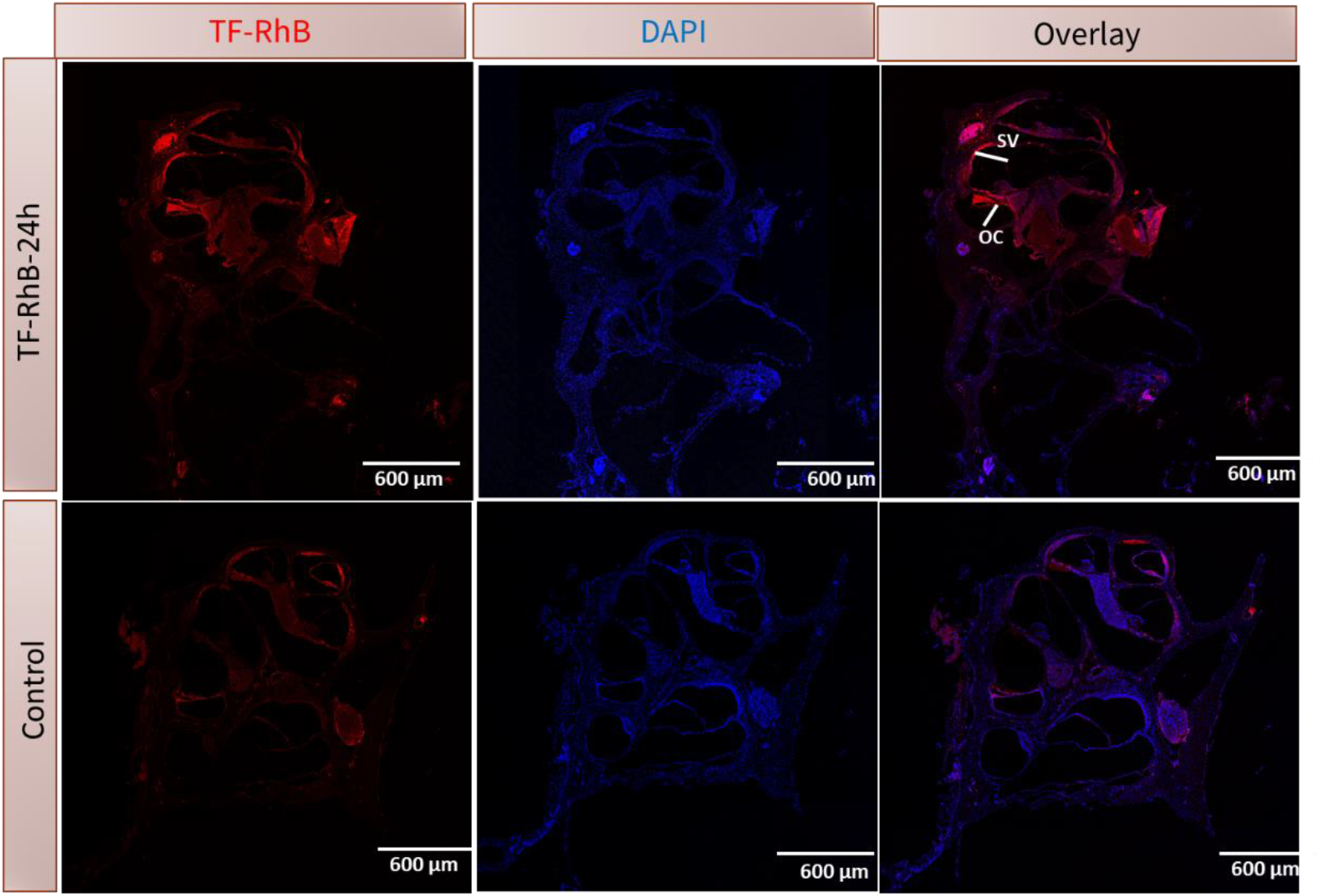
Confocal microscopy image of inner ear after 24 h of TLipo-RhB administration. DAPI-stained cell nuclei in blue and TLipo-RhB are seen in red. Abbreviations: stria vascularis (SV); organ of corti (OC) (n=5)

**Figure 6.**
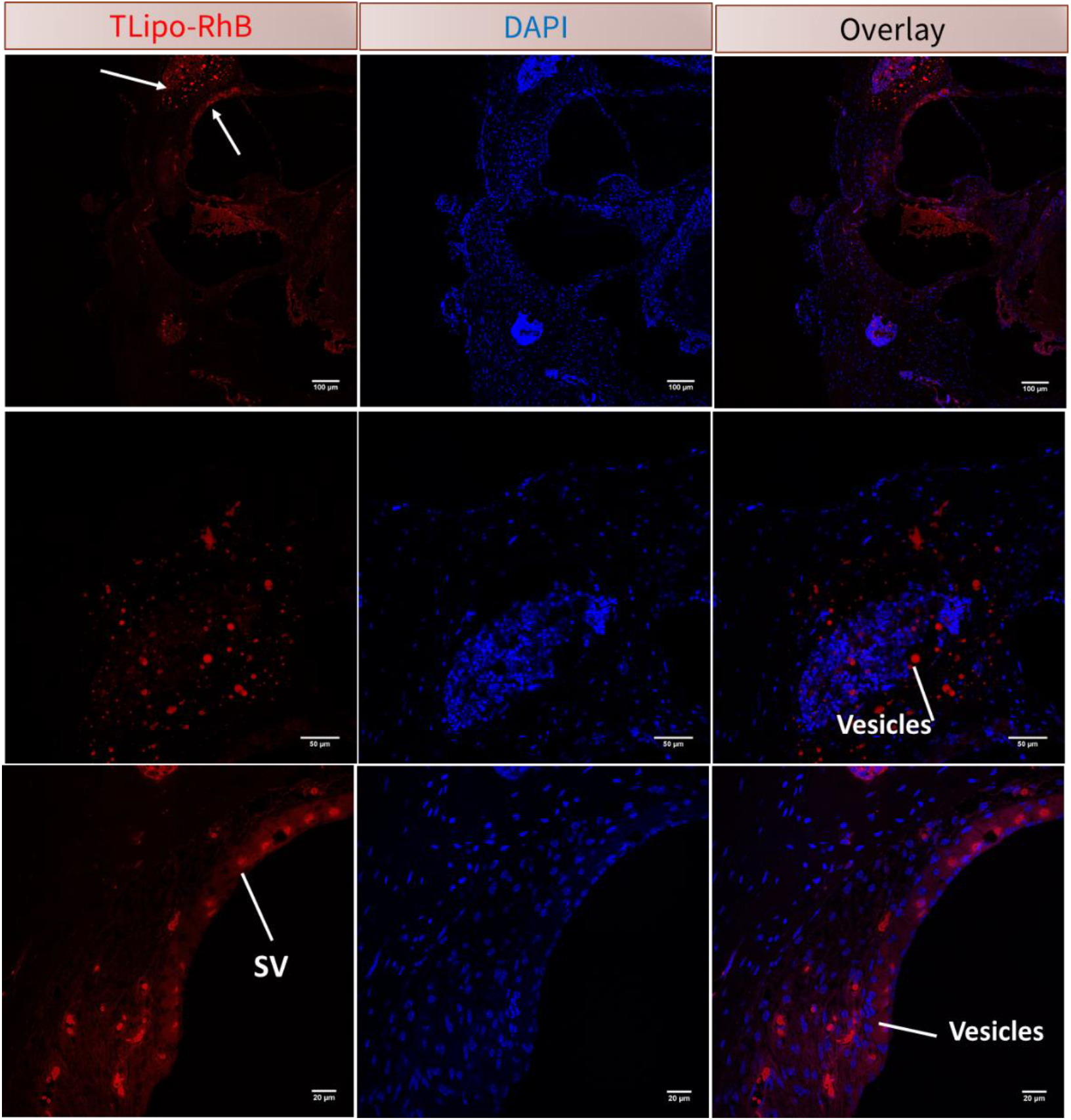
High magnification Confocal microscopy image of inner ear after 24 h of treatment. DAPI-stained cell nuclei in blue and red represents TLipo-RhB vesicles. Abbreviation: SV, Stria Vascularis; OC, Organ of Corti (n=5)

Vibratome sectioning, followed by immunohistochemistry and confocal imaging, allows for the visualization of the organ of corti. We measured the vesicles uptake in the cochlea and monitored the hair cells and the microarchitecture of the organ of corti. Neither the hair cells nor the organ of corti were damaged after 1 week of exposure. Confocal images taken from internal regions in the vibratome section showed excellent preservation of cellular morphology that is limited only by fixation artifacts (Figure 7).

**Figure 7.**
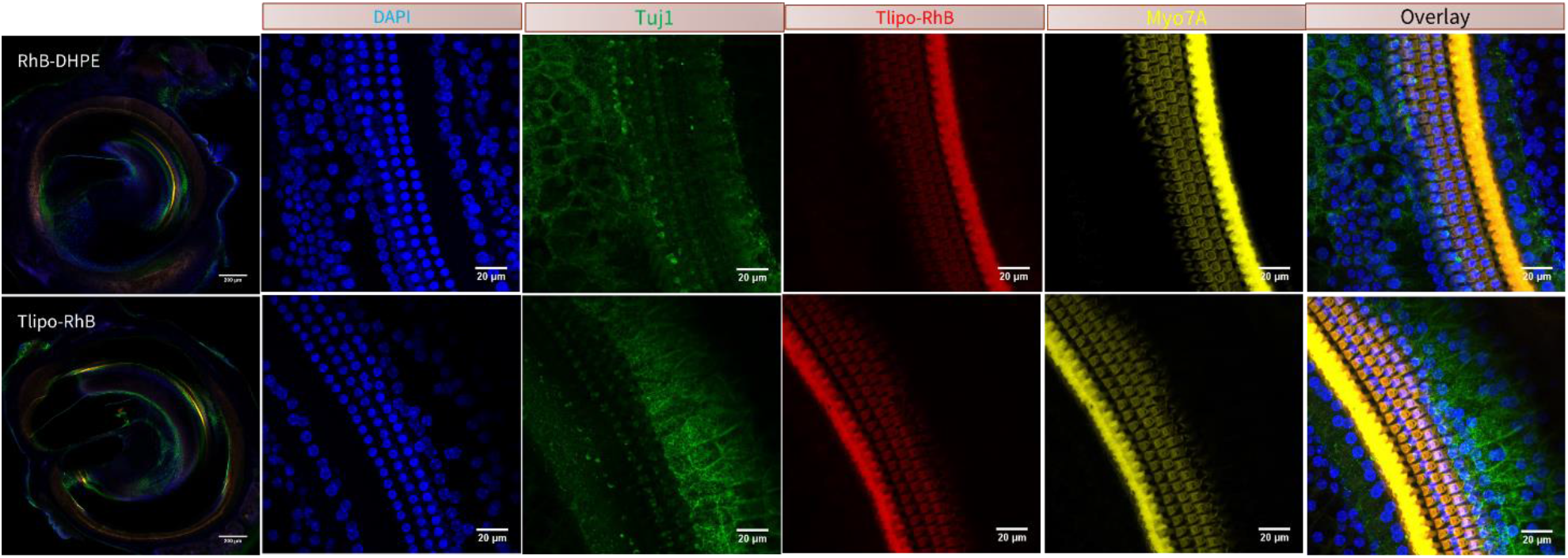
Confocal microscopy image of cochlear after 24 h of treatment. The slices were immunolabeled with Myosin 7A (Myo7A) for stains hair cells (yellow) and class III beta-tubulin (Tuj1) for neuronal structures (green) (n=5).

### TLipo has no ototoxicity

We used ABR to assess cochlear function. The ABR results showed that hearing sensitivities remained largely unchanged after TLipo administration, demonstrating that vesicles neither disturb the function of the inner ear nor cause hearing impairment in mice overtime (Figure 8a-d). To determine if TLipo vesicles caused any tissue toxicity, we assessed histological changes in the inner ear. For histology analysis, we stained the cochlea from TLipo-treated mice using hematoxylin and eosin and compared with mice without any treatment (control). No adverse effect or tissue damage were observed due to administration of TLipo vesicles. The substructure including the size of endolymph, stria vascularis, and spial ligament, the shape of spiral limbus, the number of spiral ganglion neuron, and the organ of corti did not change significantly after administration of TLipo vesicles (Figure 9).

**Figure 8.**
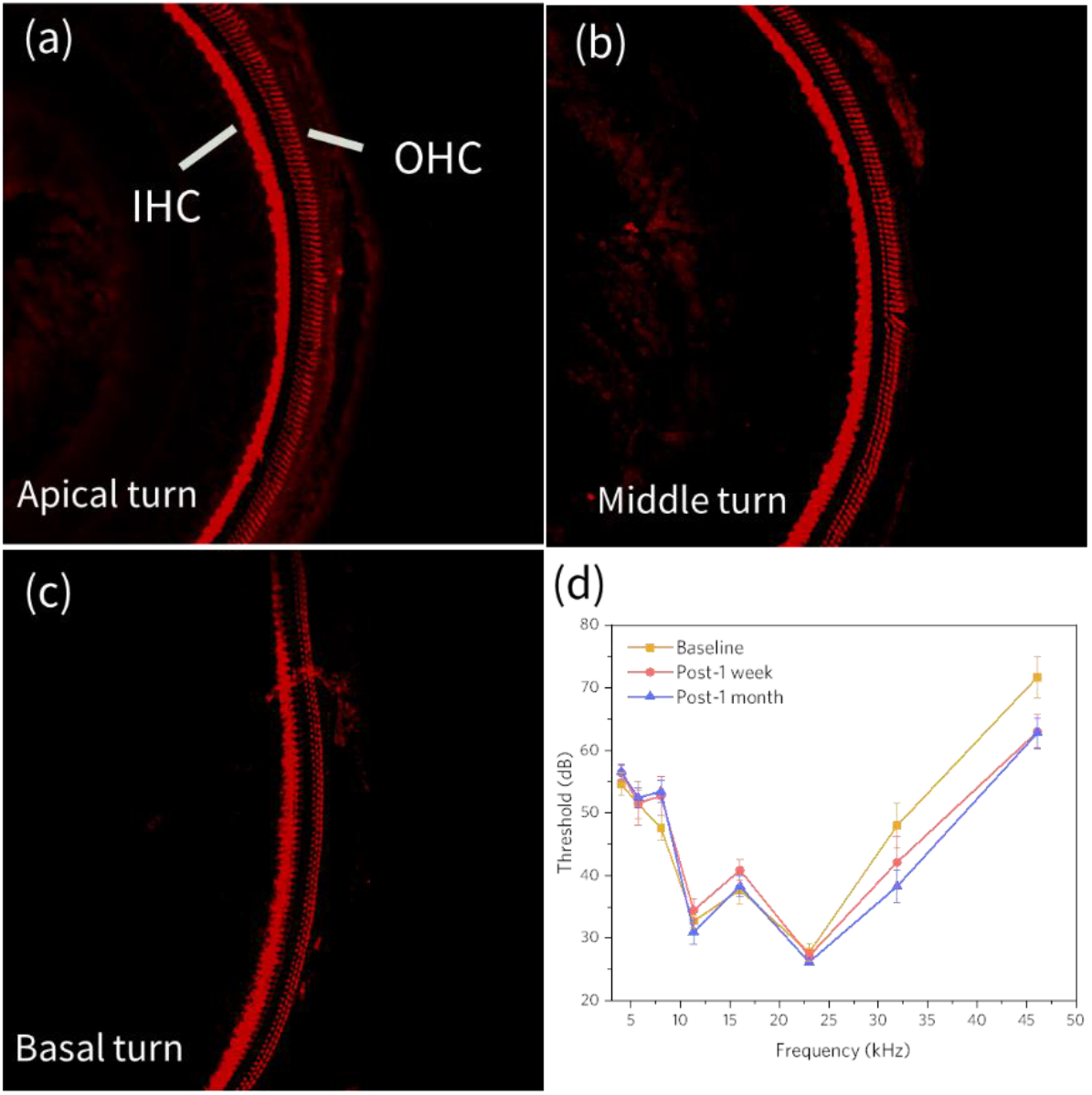
(a-c) MyosinVIIa immunostaining revealed no hair cells loss after 1 month of TLipo administration at cochlear base, middle and apex. (d) ABR thresholds of PBS (negative, baseline) and transfersome groups (n=6 in each group) after 1 week and 1 month of treatment.

**Figure 9.**
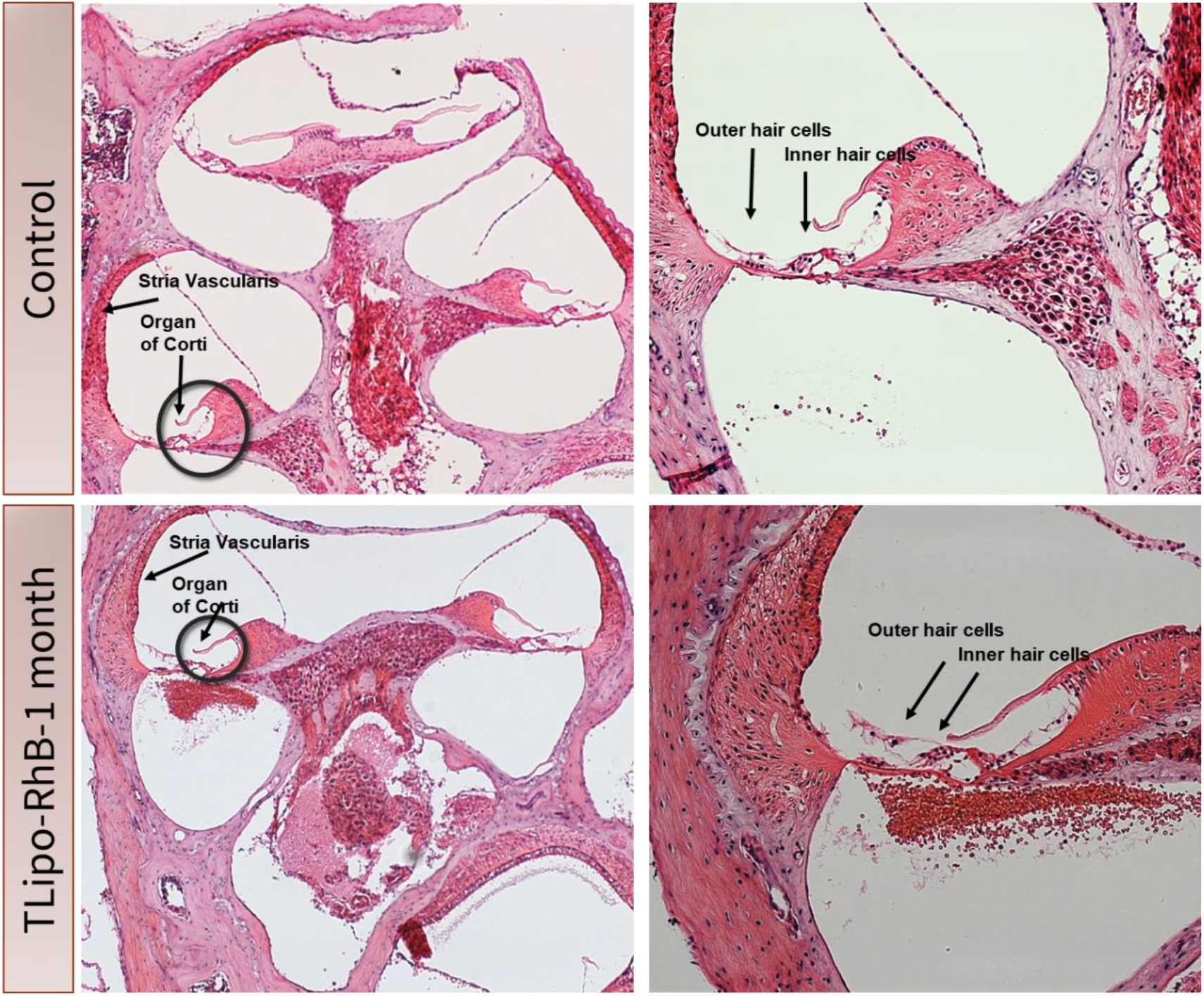
Hematoxylin-eosin staining of mouse cochleae with and without exposing to TLipo show no substructural changes after 1 month of treatment (n=5).

## Discussion

We used a subclass of liposomes called transfersomes (Scheme 1) for topical delivery of various drugs. Transfersomes are equipped with permeation enhancers which destabilize their lipid bilayers and increase their deformability. Hence, the transfersomes squeeze through the channels that can be one-tenth smaller in diameter, in lipid-rich biological barriers such as the skin or the TM. We synthesized transfersomes by the lipid film hydration method and labeled them with a Rhodamine-B (RhB) based fluorescent dye to verify their accumulation in the middle and inner ear by confocal imaging. We first verified the non-toxicity of the vesicles via in vitro and in vivo experiments. We were able to safely use the TLipo formulations at a concentration of 15.62 μg/mL while still maintaining 90% cell viability (figure 2). We then studied the penetration rate and distribution of the labeled vesicles into the middle and inner ear at different time points of the drop application by using confocal microscopy imaging. The confocal images showed that RhB-Tlipo penetrated the TM and eventually reach the middle ear after just 6 hours of dropping the formulations into the ear canal. At 1 month of post treatment, the RhB-TLipo were cleared out of the ear, suggesting that there is no accumulation (figure 3). We identified the presence of the RhB-TLipo in the inner ear with some accumulation at the stria vascularis and organ of corti (figure 6). To reach the inner ear medications must across the middle ear space to access the round window membrane (RWM) or oval window [24]. Of these structures RWM connects the middle ear and inner ear and is the most accessible entryway for inner ear drug delivery [18]. Previous studies have shown that absorption of drugs through the RWM is dependent upon permeability properties and the contact duration of the applied drug substances [26, 27]. Therefore, a variety of techniques to deliver medications to the including invasive modalities such as Silverstein Microwick® [28] and the Microcather (μCat®) [29] inner ear such as have been described. The TM presents a barrier to delivering treatment agents into the middle ear and is composed of three layers. Stratum corneum (SC) is the outer-most layer of TM and is the main barrier for drug permeation [11]. The SC is formed of highly keratinized dead cells (corneocytes) which are densely packed within the extracellular lipid matrix. There are two mechanisms for transporting the medication through the SC: 1) crossing the lipid matrix (intercellular path), and 2) using corneocytes (intracellular) [30]. To overcome these barriers, it is crucial for the agents to be highly hydrophobic [11], which is the case with our TLipo formulation.

**Scheme 1.**
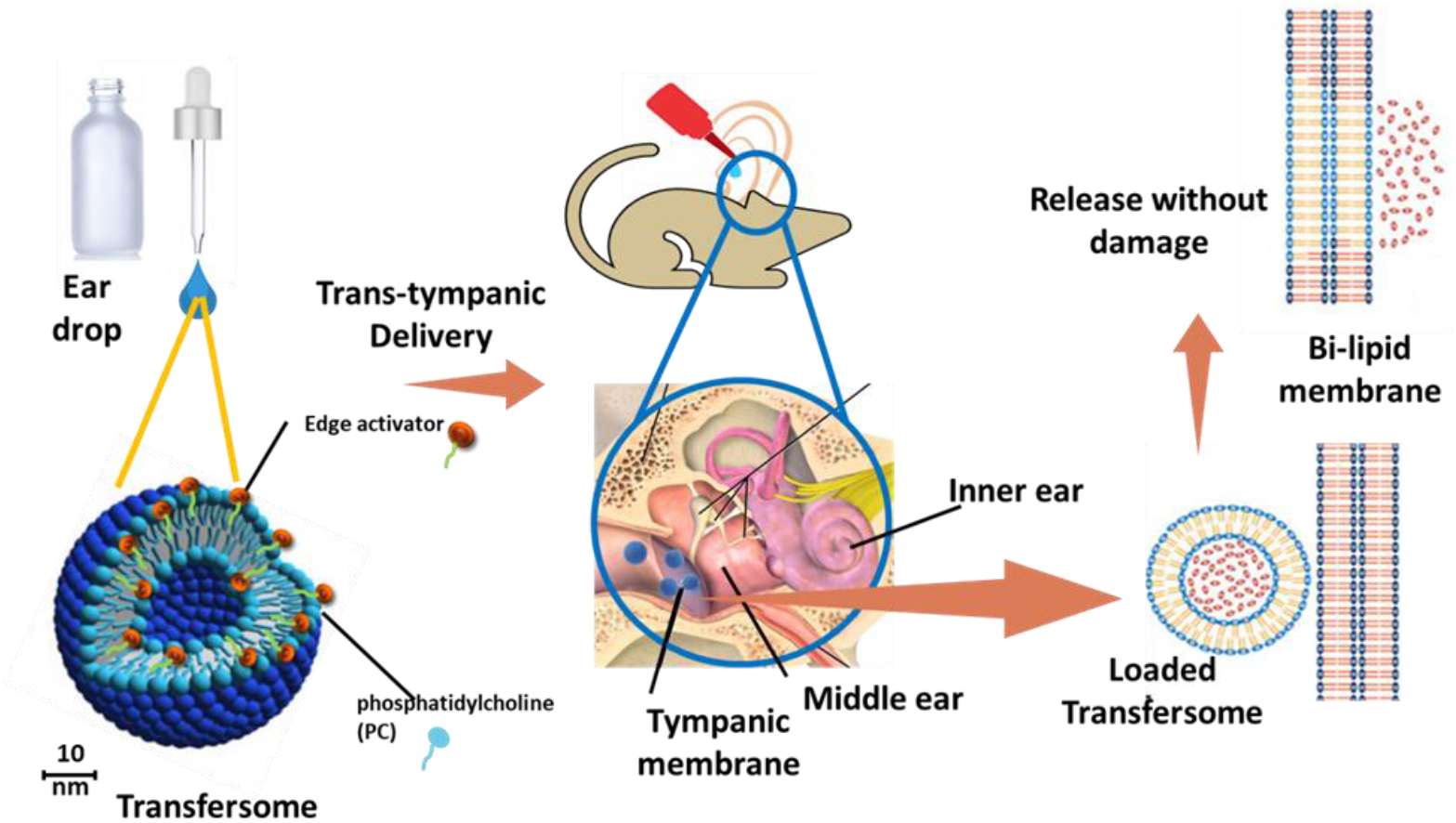
Concept for non-invasive delivery to middle ear using TLipo vesicles. The eardrop carries TLipo vesicles and penetrates the tympanic membrane without damaging it due to similar lipid structure.

This finding opens up opportunities for drug delivery to prevent or treat hearing loss. In terms of functionality, ABR thresholds for the baseline, 1 week and 1 month post-formulation drops show no significant difference, indicating that the TLipo is not toxic to hair cells at a higher concentration. In terms of structure, our histological analysis and immunohistochemistry images showed preservation of hair cells, and the organ of corti.

## Conclusion

Results of the present study showed that our TLipo formulation is an effective and safe way to deliver diagnostic and therapeutic agents to middle and inner ear. In vitro cell viability and in vivo auditory brainstem response studies in mice models indicated the safety of the TLipo delivery system. The confocal microscopy images of middle and inner ear confirmed penetration and localization of the TLipo in middle ear in 3-6 hours and their clearance from the ear in less than 1 month, making them promising imaging and drug delivery systems to diagnose and treat ear diseases. TLipo delivery can potentially provide a convenient, inexpensive, and risk-free alternative to surgery, resulting in fewer doctor visits, reducing cost of care, and improving overall quality of life for children and their parents. However, extensive pe-clinical studies are required before TLipo can be verified as a viable solution. Future studies will start this process by evaluating encapsulation of therapeutic drugs for topical treatment of middle ear conditions.

## Acknowledgment

This work was supported by the Stanford Predictives and Diagnostics Accelerator (SPADA) program. The authors are thankful to Sara Talaee for assisting with 3D visualization of the cellular confocal images using Imaris, Elizabeth Montabana for Cryo-TEM images (Stanford SLAC CryoEM Initiaitve), and Doreen Wu for H&E staining.

